# Large scale semi-automated quantification of viral spread reveals multi-phasic structure of plaque formation

**DOI:** 10.1101/2024.11.18.624223

**Authors:** Liam M. Howell, Timothy P Newsome

**Affiliations:** School of Life and Environmental Science, The University of Sydney, Australia; Sydney Institute for Infectious Diseases, The University of Sydney, Australia

**Keywords:** fluorescence microscopy, image analysis, poxvirus, virus replication and spread, plaque assay

## Abstract

Plaque assays are typically quantified at an endpoint and fail to capture the temporal dynamics of viral infection and spread. In this study we introduce a high-throughput method utilising live-cell imaging and a recombinant vaccinia virus (VACV) engineered to express two fluorescent transgenes linked to distinct replication phases. This novel approach allows for real-time tracking of VACV plaques from their inception at a single infected cell through to multicellular expansion, enabling detailed kinetic analysis of plaque development. Using this analysis pipeline we categorise VACV plaque formation into three distinct phases: Establishment, Expansion, and Exhaustion, and quantitatively describe their dynamics. Our findings reveal significant variability in the initiation time of plaque spread (from single-cell to multicellular stages) and the growth rate of plaques, both between individual plaques and across different cell lines. This variability underscores the influence of host cellular factors on the kinetics of viral replication. Additionally, we reveal that the viral replication cycle dramatically accelerates through the early phases of plaque formation, which we hypothesise is driven by the changing multiplicity of cellular infection at the plaque edge. Finally, we demonstrate that differences in plaque sizes between two cell lines (BS-C-1 and HaCaT) can be largely attributed to variations in viral replication rate. This research reinforces the value of live-cell fluorescence microscopy in elucidating the complex spatiotemporal dynamics of viral infections, and contributes to a deeper understanding of the mechanisms driving viral spread and replication kinetics.

## Introduction

The plaque assay is a core virological technique for the enumeration of infectious units of viruses that induce a cytopathic effect. A susceptible cell monolayer is infected with virus and the free diffusion of progeny virions is inhibited with a semi-solid or solid overlay. Cells are incubated over a suitable time course allowing for multiple rounds of replication and cell-to-cell transmission. Finally, quantitation of infectious units by the detection of zones of cytopathic effect, often through the use of a cell stain, is performed [1]. The plaque assay is exquisitely sensitive to small changes in replication kinetics and is capable of discerning subtle phenotypes due to the amplifying effect of multiple replication cycles.

In its canonical form, the plaque assay provides a static, endpoint quantification of infectious units and an assessment of virus spread, but does not capture the temporal dynamics of this process. Recent applications of live-cell imaging are bridging this gap, but have been primarily focused on reducing assay run time, subjectivity and ambiguity, and increasing sensitivity [2–4]. Although there is a considerable body of work centred on modelling the kinetics of cell-to-cell virus spread, direct, high temporal resolution imaging and analysis of this critical process is lacking [5, 6]. The integration of time-lapse fluorescence microscopy, especially with recombinant viruses expressing distinct fluorescent markers for different stages of viral gene expression [7], offers a powerful means to examine the dynamics of plaque formation and the kinetics of viral replication within a plaque, greatly enhancing our understanding of the spatiotemporal dynamics of viral propagation.

In this study, we utilise live-cell imaging with a fluorescent-reporter-expressing vaccinia virus (VACV^Rep^) to investigate viral spread and replication dynamics at high temporal resolution. This approach enabled distinct visualisation of early and late phase viral gene expression. We observed the development of viral plaques from a single infected cell to multicellular foci and characterised the kinetics of viral spread. By tracking the spatiotemporal dynamics of these two reporters we were able to capture variation in the replication cycle duration throughout the lifetime of a plaque. Our analysis delineated plaque formation and expansion into three phases: establishment, linear expansion, and a growth exhaustion phase. Additionally, we explored variations in plaque size across different cell lines, identifying significant differences in the timing of plaque spread initiation and growth rates. These findings quantify the influence of cell line-specific factors on viral replication and spread kinetics. This study demonstrates the utility of real-time imaging in understanding viral infection dynamics and highlights the complexities of plaque formation and viral spread in different cellular environments.

## Results

### Tracking viral plaque spread by live-cell time-lapse fluorescence microscopy

To monitor the stages of infection in real-time, we engineered a VACV reporter virus by recombining viruses bearing the fluorescent markers pE/L-mCherry [8] for Early/Late and pA3L-YFP-A3 [9] for Late gene expression. The resulting recombinant, termed VACV^Rep^, was isolated by multiple rounds of plaque purification. The use of VACV^Rep^ allowed for identification of infection through mCherry fluorescence and facilitated tracking of late gene expression via YFP fluorescence (Fig. 1A).

**Figure 1:**
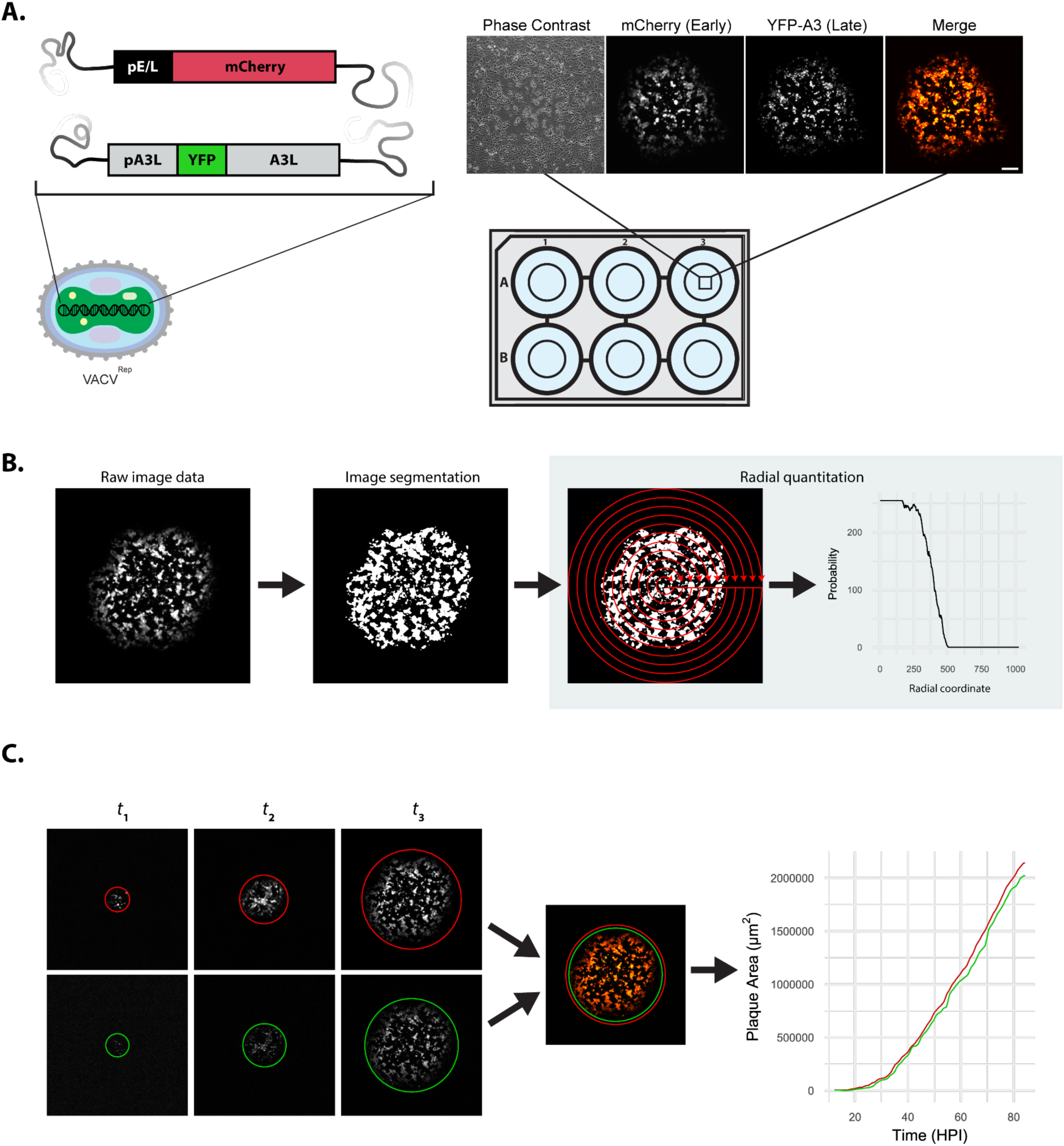
Capturing VACV plaque growth dynamics. **A**, Schematic and micrographs depicting VACV^Rep^ infection of BS-C-1 cells, demonstrating Early (mCherry) and Late (YFP) fluorescent reporter expression. The schematic illustrates the transgenes used for reporter expression, while the micrographs display the reporter fluorescence pattern within a plaque, captured through phase contrast and fluorescence microscopy. **B**, VACV^Rep^ plaque image analysis workflow. A trained random forest classifier processes raw fluorescence micrographs and converts pixel values from integrated densities to probabilities, distinguishing between infected and uninfected regions of each image. Radial quantitation then translates these probability maps into radial profiles, representing the probability that the plaque edge is found at a given distance from the centre of the image. **C**, Applying the image segmentation and radial quantitation approach to sequential time-lapse images captures the progressive expansion of the plaque, visualised by both Early and Late reporter expression. Scale bar: 400 µm (A).

To capture the dynamics of viral plaque spread, a confluent monolayer of BS-C-1 cells was infected with VACV^Rep^ at an MOI sufficient to obtain well separated plaques (20 PFU/well in a 6-well plate), and imaged by fluorescence microscopy and phase contrast at 10-minute intervals over a 72-hour period. As a result of the exceptionally early expression of mCherry we were able to identify and begin imaging single infected cells (putative plaques) as early as 8 hours post infection (HPI). This method provided a detailed visualisation of plaque development and spread within the cell monolayer (Fig. 1A). The fluorescence channel images were then subjected to machine learning-augmented segmentation to identify infected cells, converting pixel values from intensities to probabilities (Fig. 1B). These segmented images were converted to radial profiles, representing the probability of the plaque edge being located at varying radii from the centre of the image. Applying this segmentation and radial profiling process to all frames of the time-series data enabled accurate tracking of viral plaque growth over time (Fig. 1C, Supplementary Video 1).

### Analysing the dynamics of VACV plaque growth

The growth kinetics of VACV^Rep^ plaques were characterised by readily apparent variation in both the time required for progression from a single-cell infection to a multicellular plaque and the rate of plaque expansion. Time-course analysis of plaque radius and area showed a consistent growth velocity in each plaque until approximately 60 HPI, followed by a phase of deceleration (Fig. 2A, B).

**Figure 2:**
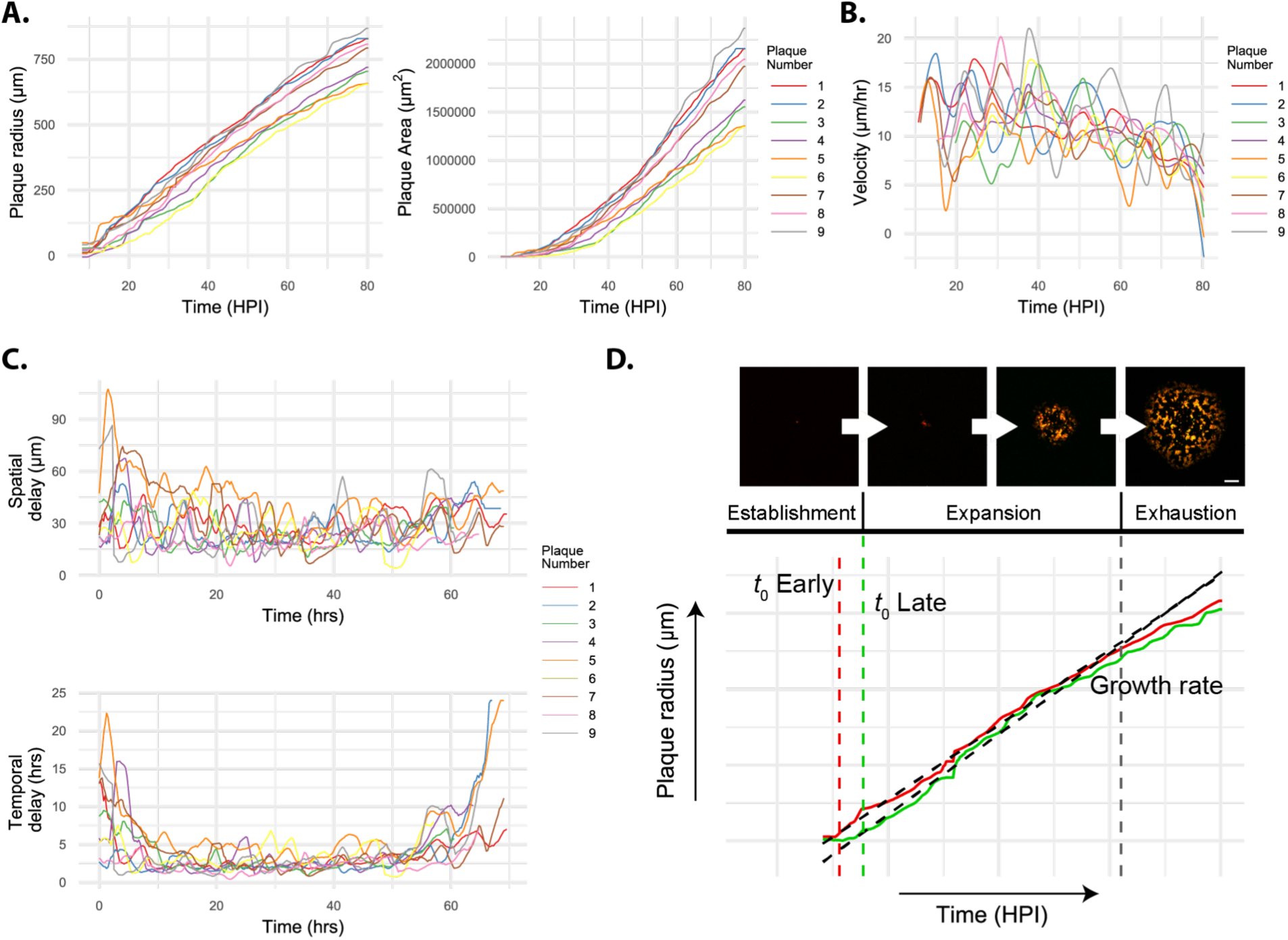
Modelling VACV plaque growth dynamics. **A**, Graphical representation of the radius and area of multiple VACV^Rep^ plaques over time as determined by Early reporter expression. **B**, Rate of VACV^Rep^ plaque expansion over time. **C**, Analysis of the spatial and temporal interval between Early and Late reporter edges for multiple plaques, representing the delay between early and late gene expression in infected cells over time. This delay serves as an indicator of the rate at which the VACV^Rep^ replication cycle progresses in individual cells at the edge of the plaque as it expands. **D**, Decomposition of plaque expansion into distinct phases: the establishment phase, characterised by a single infected cell signifying plaque initiation; the expansion phase, where the plaque grows at a steady rate modelled linearly; and the exhaustion phase, where growth decelerates. This model elucidates the full progression of plaque development. Scale bar represents 200µm.

Spatial analysis of the plaques revealed a clear separation between the furthest extent of Early and Late reporter expression at the periphery of plaques. By measuring the distance and velocity of these two regions, we inferred the temporal lag in early versus late gene expression at the edge of a plaque. This approach provided insights into the viral replication rate at the plaque periphery over time. Our data indicated a rapid increase in replication rate in newly infected cells during the first 10-15 hours of plaque development. This initial phase was succeeded by a sustained plateau and subsequently by a deceleration phase beyond approximately 60 HPI (Fig. 2C).

Based on these data, we propose that plaque formation occurs in three distinct phases: (1) the initial establishment phase in one cell followed by spread to multiple cells; (2) a linear expansion phase characterised by a constant growth rate; and (3) an exhaustion phase where the growth rate declines due to the depletion of available resources or the accumulation of cell population defences (Fig. 2D). Employing changepoint analysis and linear regression, we automated the detection of key temporal events: the initial spread of infection (*t*_0_ Early), the commencement of the second replication cycle within the plaque (*t*_0_ Late), and the plaque’s growth rate during its expansion phase.

### Differential plaque spread dynamics in epithelial cell lines

Variability in viral plaque size across different cell lines is a well-documented phenomenon [10], yet the underlying dynamics that contribute to this variability remain poorly understood. To investigate this, we selected two cell lines: BS-C-1, a line of African green monkey kidney cells commonly used for the cultivation of viruses [11], and HaCaT, an immortalised human keratinocyte line that provides a model for human skin [12–14], a common target of poxvirus infection [15, 16].

Analysis of plaques in both cell lines showed that plaques in BS-C-1 cells typically reached a larger size than those in HaCaT cells (Fig. 3A). When examining the onset of plaque spread, BS-C-1 cells displayed a significantly later start (*t*_0_) than HaCaT cells (Fig. 3B) and, conversely, a significantly faster growth rate (Fig. 3C, Supplementary table 1, Supplementary table 2). Notably, there was increased inter-plaque variability in the timing of spread initiation (*t*_0_) in BS-C-1 cells, whereas HaCaT cells showed more variation in growth rate. This difference in the initiation and rate of spread likely represent the contribution of cell line-specific factors that influence viral replication kinetics.

**Figure 3:**
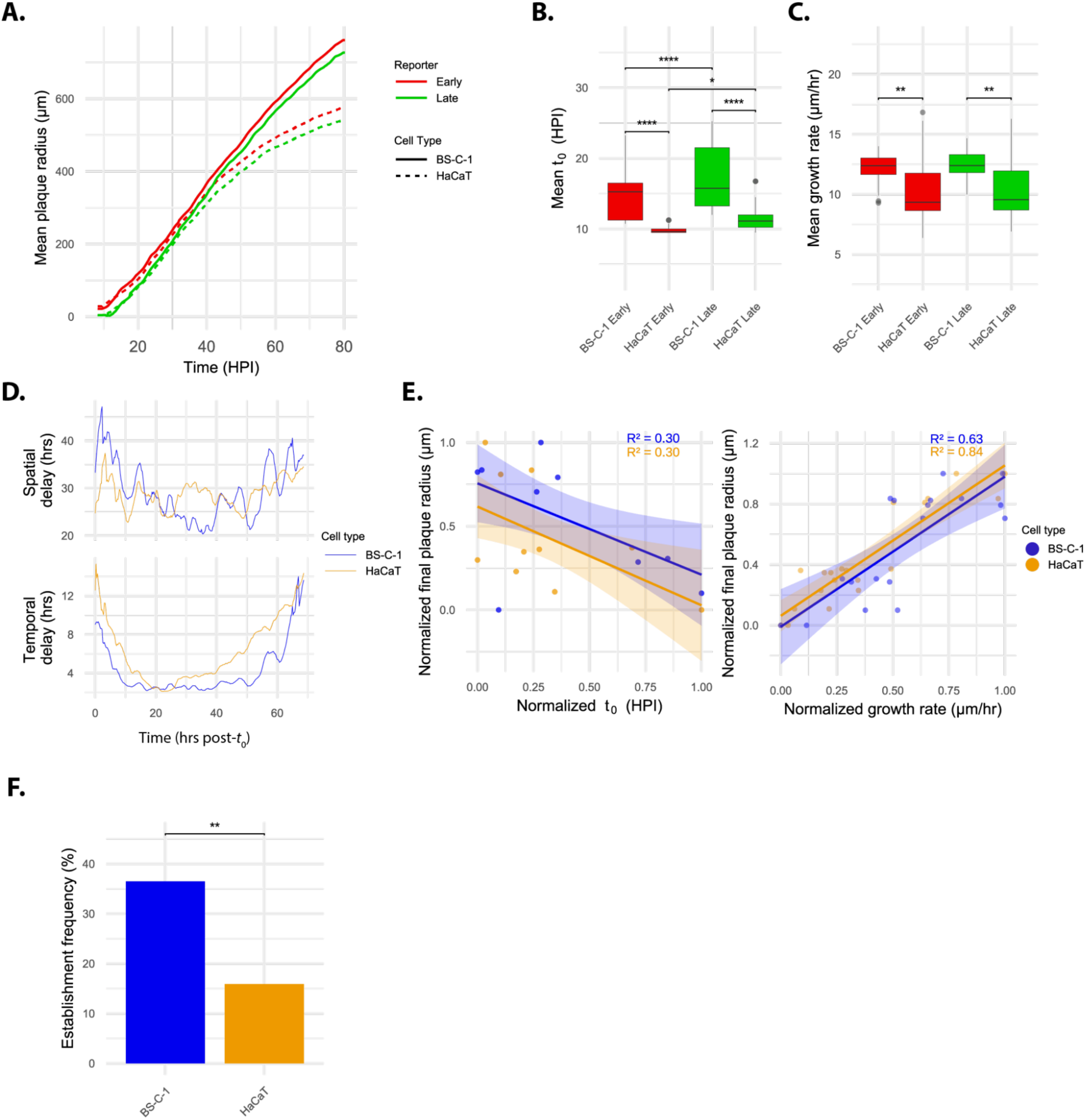
Differential VACV plaque dynamics between epithelial cell types. **A**, The mean radius of VACV^Rep^ plaques in BS-C-1 or HaCaT cells over time, using Early and Late reporters to track growth (number of plaques: HaCaT; n = 10, BS-C-1; n = 9). **B**, **C**, Comparison of the time at which plaque spread begins (*t*_0_) and the rate of plaque growth between BS-C-1 and HaCaT cells. **D**, The average spatial and temporal delay between Early and Late reporters within BS-C-1 and HaCaT cells. Plaques were synchronised by their respective *t*_0_ values prior to mean calculations. **E**, Correlation between the final radius of plaques and *t*_0_, and the correlation between the final plaque radius and the growth rate during expansion (right), for both BS-C-1 and HaCaT cells. Correlation strength is indicated by R² values, with linear regression lines plotted over the data points, and the 95% confidence interval shown as reduced opacity bands around the regression lines. Features are scaled for each cell type to enable direct comparisons. **F**, Comparison of the percentage of single infected cells that successfully establish plaques in BS-C-1 versus HaCaT cell lines, providing a measure of the relative efficiency of plaque formation between the two cell types. Statistical significance for all comparisons was determined using pairwise T-tests with Bonferroni’s correction to adjust for multiple comparisons where appropriate.

By aligning the time series data from each plaque according to their respective *t*_0_ and examining the average dynamics of the relationship between Early and Late reporters, we observed that viral replication within plaques initially accelerated, plateaued for a period, and then decelerated in both cell lines (Fig. 3D). However, the plateau phase — where plaques expanded at their maximum rate — was sustained for longer in BS-C-1 cells.

Given the observed variability in *t*_0_ and growth rate, we explored their relationship with final plaque size using linear regression (Fig. 3E). Growth rate was found to be a greater determinant of final plaque size than *t*_0_, explaining 63% and 84% of the variability in BS-C-1 and HaCaT cells, respectively. A multivariate regression model incorporating both *t*_0_ and growth rate accounted for the majority of variance in final plaque size in both cell lines. In BS-C-1 cells, the model was highly predictive (R² = 0.79), with *t*_0_ (p = 0.00484) and growth rate (p = 3.21 x 10^-5^) both significantly contributing to final plaque size.

For HaCaT cells, the model also demonstrated strong predictive ability (R² = 0.85), but only the growth rate (p = 3.49 x 10^-7^) was a significant factor, with *t*_0_ not having a significant impact (Fig. 3E).

By imaging a large number of single infected cells and monitoring their transition to plaques, we were also able to observe the frequency at which this transition occurs successfully. The successful transition from a single infected cell to a mature plaque is a critical parameter in understanding viral infection, as it encapsulates the efficacy of early viral replication and cell-to-cell spread in an environment of high virus-host competition. The probability of a single infected cell progressing to a plaque was significantly greater in BS-C-1 cells compared to HaCaT cells (Fig. 3F). Interestingly, both cell lines produced a similar total number of plaques (data not shown) when infected at the same MOI (titre determined by plaque assay in BS-C-1 cells).

## Discussion

In this study, we offer a detailed analysis of the temporal dynamics of plaque formation by a large DNA virus. Plaque assays, a cornerstone technique in virology, are a key method for assaying virus spread and replication. The dimensions of a plaque reflect the cumulative impact of numerous viral replication cycles, thus serving as an exceptionally sensitive gauge for assessing viral attenuation across diverse replication and spread-related factors. By combining real-time imaging and a novel fluorescent reporter virus with our semi-automated analysis of plaque formation we extended the utility of this assay, extracting a wealth of additional information and enabling the detection and quantification of a wider spectrum of viral phenotypes and nuances in virus-host interactions.

We tracked the spread of a large number of early infection events from single-cells to late-stage plaques and delineated viral spread into three distinct phases: Establishment, Expansion, and Exhaustion. Employing linear modelling and changepoint analysis, we established a semi-automated methodology to demarcate these phases. This method allowed us to quantify critical metrics of plaque formation and spread, including the initiation of multicellular infection (*t*_0_) and the rate of plaque expansion.

Our findings reveal high levels of variability in the time taken for infections to spread from a single cell, and in the rate at which they spread after passing this key bottleneck. In BSC-1 cells, the average growth rate during the Expansion phase was observed to be approximately 12 µm/hr (Supplementary table 1). With the average cell diameter being 38.64 ± 2.27 µm (SEM, n= 48), we estimate a minimum cell-to-cell spread time of 3.2 hours, implying a VACV replication rate markedly more rapid than typically reported [17, 18]. Intriguingly, we were able to detect variation over time in the temporal delay between early and late viral gene expression at the edge of plaques, consistent with a minimum of 2.1 hours from early gene expression to progeny virion production. These findings suggest that accelerated replication kinetics during the expansion phase substantially account for the rate of spread, complemented by VACV-induced cell motility [19, 20], challenging previous assumptions made in considering models of VACV cell-to-cell spread [21, 22]. Additionally, single-cell VACV^Rep^ data we have collected profiling infection dynamics in HeLa cells (data not shown) indicate that the delay between early and late gene expression correlates with MOI. Although reliant on a number of assumptions, this suggests that a dramatic variation in local MOI at the plaque edge occurs (from approximately 1, in the initial spread to multiple cells, to 74 at the peak rate of expansion), and that this greatly influences the duration of the replication cycle at the single cell level.

In line with existing research, we observed variations in plaque size across different cell lines, likely influenced by varying susceptibility to infection and the robustness of innate immune responses [20, 23–25]. Our comparative study of BS-C-1 cells and HaCaT keratinocytes revealed distinct differences in plaque growth dynamics. BS-C-1 cells demonstrated a delayed *t*_0_ but a prolonged Expansion phase, indicating a higher resistance to initial infection and a greater capacity for sustained viral replication compared to HaCaT cells. This difference could be attributed to the differences in innate immune induction, as keratinocytes exhibit a diverse array of antiviral defences and potent innate immune activation capabilities [26–29]. Furthermore, there is evidence of IFN-β and IFN-λ mediated arrest of viral spread in HaCaTs, despite replication proceeding in single cell infections [30–32], which may explain the reduced Expansion phase seen in our data. Interestingly, we found that VACV infection spread through a HaCaT cell monolayer at a maximum rate of 1.9 hours per cell, based on an average growth rate of 10.4 µm/hr (Supplementary table 2) and an average cell diameter of 20.40 ± 0.61 µm (SEM, n= 48). This indicates a rate of cell-to-cell transmission considerably faster than that seen in BS-C-1 cells. However, both cell types exhibited a similar minimum Early-Late temporal delay (around 2.1 hours). The faster spread in HaCaT cells likely reflects their stronger migratory ability, enabling quicker expansion of the plaque boundary compared to the time taken for virus to spread between adjacent cells [19, 33].

In analysing the relationship between *t*_0_, growth rate and final plaque size, we found that these two features accounted for the majority of variation. However, growth rate was a less important determinant in BS-C-1 cells, consistent with our observations of a reduced Expansion phase duration in HaCaTs. This highlights the critical role of the timing of infection initiation and viral spread rate in determining the extent of infection across different cell types.

Our integration of real-time imaging, fluorescent reporter viruses, and a simple model of plaque expansion has yielded a more nuanced understanding of the intricate dynamics of virus-host interactions that drive cell-to-cell spread. This study not only enhances our knowledge of viral replication and spread mechanisms but also offers valuable insights that could be applied to developing effective antiviral strategies and improving quantification of viral phenotypes.

## Methods

### Cell culture

All cell lines were cultured in Dulbecco’s Modified Eagle’s Medium (DMEM, Gibco) supplemented with 10% foetal bovine serum (FBS, Australian origin, Bovogen) for HaCaT cells (ATCC, CCL-2) and 5% for BS-C-1 cells (ATCC, CCL-26). Additionally, 1% PSG (Gibco), containing 100 units/mL penicillin, 100 µg/ml streptomycin, and 292 µg/ml L-glutamine, was added. Cells were incubated at 37°C with 5% CO2 in a humidified environment. Regular testing was conducted using the MycoAlert® Mycoplasma Detection Kit (Lonza Bioscience) to ensure the absence of Mycoplasma contamination.

### Plaque assays

BS-C-1 or HaCaT cells were seeded into 6-well tissue culture plates (SPL Life Sciences, 6-well plate) and grown to confluence. Virus strains were diluted in SFM to the appropriate concentration and added to cells, after first aspirating growth medium and washing cells once with PBS. Cells were incubated at 37°C, 5% CO_2_ for 1 hr, washed again with PBS, then overlaid with appropriate medium. For live imaging cells were overlaid with 1x Minimal Essential Medium (MEM, Gibco) containing 1.5% carboxy-methyl cellulose (CMC) and supplemented with 1% PSG (Gibco, 100 units/mL of penicillin, 100 µg/ml of streptomycin and 292 µg/ml L-glutamine). For plaque isolation cells were overlaid with 1x Minimal Essential Medium (MEM, Gibco) containing 0.9% (w/v) Ultrapure Agarose (Invitrogen) and supplemented with 1% PSG (Gibco, 100 units/mL of penicillin, 100 µg/ml of streptomycin and 292 µg/ml L-glutamine). Both media variants were supplemented with 10% FBS for HaCaT growth or 5% FBS for BS-C-1 growth.

### Construction and characterisation of VACV^Rep^

VACV^Rep^ was generated by recombination of VACV WR pE/L-mCh and VACV WR YFP-A3 in BS-C-1 cells co-infected for 48 hrs. Post-infection, cells were harvested and lysed via freeze-thaw cycles. The lysates underwent serial dilution and were used to infect confluent BS-C-1 cell monolayers. Plaques exhibiting both pE/L-mCh and YFP-A3 fluorescence were identified, isolated, and underwent six rounds of plaque purification on BS-C-1 cell monolayers.

### Virus stock preparation

Virus stocks were purified from the cytoplasmic fraction of infected cells by zonal sucrose gradient centrifugation (as in [11]). Titres of all virus stocks were determined by plaque assays on BS-C-1 cells.

### Live fluorescence microscopy of VACV^Rep^ plaques

Cells were cultured until confluent then infected with VACV^Rep^, followed by overlaying with 1x MEM containing CMC (as described above under ‘Plaque assays’). At approximately 8 HPI, mCherry fluorescence allowed identification of individual infected cells, enabling the commencement of live cell imaging. This imaging was conducted using an Eclipse Ti-E Microscope (Nikon), equipped with a motorised stage, an iXon Ultra 888 EMCCD camera (Andor), in an environment maintained at 37°C with 5% CO_2_ (Okolab, Cage incubator).

Images were captured every 30 minutes for a duration of 72 hrs, in Phase Contrast and both red (TxRED Semrock, 500ms exposure) and green (FITC Semrock, 500ms exposure) channels, with a 4x objective. NIS-Elements Microscope Imaging Software AR (Nikon, v4.51.01) was employed for image acquisition control.

### Image analysis

Initial segmentation of raw fluorescence channel images was conducted using a trained random forest classifier in Ilastik [34] to convert raw fluorescence intensity images to probability maps. This process substantially reduced noise and artefacts in the raw fluorescence channel data.

Subsequently, a custom Fiji [35] macro integrating the Radial Profile Angle plugin [36] was utilised to transform these two-dimensional probability maps into probability profiles. This transformation involved drawing concentric circles on each image, with each circle incrementally increasing in diameter by one pixel. The macro then calculated the mean pixel intensity beneath each circle’s circumference.

Repeated across images from all timepoints, this method generated a time-resolved series of probability profiles. These profiles indicated the likelihood of each circular area in the images representing either plaque or background, with a radial resolution of one pixel. This approach provided a precise spatial and temporal analysis of the dynamics of viral spread through a monolayer of cells.

### Data analysis

The dynamics of viral plaque growth can be conceptualised as encompassing three distinct phases:

1. **Establishment phase**: This initial phase signifies the period when the single infected cell that seeds a plaque hasn’t yet infected its immediate neighbours.
2. **Expansion phase**: Following establishment, this phase reflects the time when the infection has spread to multiple cells and begins to progress outwards at a relatively constant rate.
3. **Exhaustion phase**: Following expansion, this phase represents the period where plaque expansion begins to slow. The timing (and presence) of this phase is impacted by a number of factors related to cell type and culture environment.

To automate the estimation of the time point at which the establishment phase ends and the expansion phase begins (*t*_0_) we employed a multi-phase process.

First, we calculated the rate of change in the plaque’s radius over time. This is effectively the velocity of the plaque’s growth. This step involves differentiating the data with respect to time to obtain the first derivative, which is subsequently smoothed using LOESS regression to reduce the influence of short-term fluctuations and noise. The smoothed estimated rate of change of a plaque’s radius at any given discrete time can therefore be represented by:

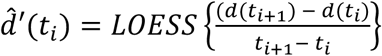

where d(*t*) is the radius of the plaque at time *t*, *t*_i_ is the current time point, and *t*_i+1_ is the subsequent time point.

Second, we applied changepoint detection algorithms (using the PELT method) [37] on the smoothed derivative data to detect local changes in both the mean and variance, identifying points in time where there is a statistically significant shift in the growth rate of the plaque. To minimise the impact of noise we implemented 3 criteria that a changepoint must satisfy to be considered valid:

1. The average plaque radius after the changepoint must exceed that before the changepoint by a predetermined factor.
2. The increase in plaque radius must be greater than a minimum absolute value.
3. There must be no negative growth rates detected in a defined period following the changepoint, as this would contradict the ground truth of the phenomenon.

The earliest time point that satisfies these criteria is then taken as *t*_0_.

With *t*_0_ identified, data between *t*_0_ and 50 HPI (a conservative estimate of the onset of the Exhaustion phase) was isolated. Within this window, a linear regression model was applied to the plaque radius as a function of time, represented as:

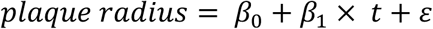

Where *t* represents a discrete time point, *β*_0_ represents the initial radius of the plaque, *β*_1_ represents the growth rate of the plaque, and ɛ denotes the error term.

### Statistical analysis

All statistical analyses were performed using R statistical software v2023.9.0.463 [38], primarily with functions implemented through the rstatix package [39]. Sample size, number of replicates, statistical tests performed, and the nature of any ɑ-value corrections applied are indicated in the appropriate figure legends. Annotations of significance indicate the following p-values: * p<0.05, ** p<0.01, ***, p<0.001, **** p<0.0001.

## Supplementary information

**Supplementary Video 1: Tracking plaque growth over time.**

A time-lapse visualisation of the output given by the automated plaque edge detection and analysis pipeline, demonstrating the simultaneous tracking of Early and Late reporter expression. Time post infection is indicated in hours:minutes. Scale bar represents 400µm.

**Supplementary Figure 1.**
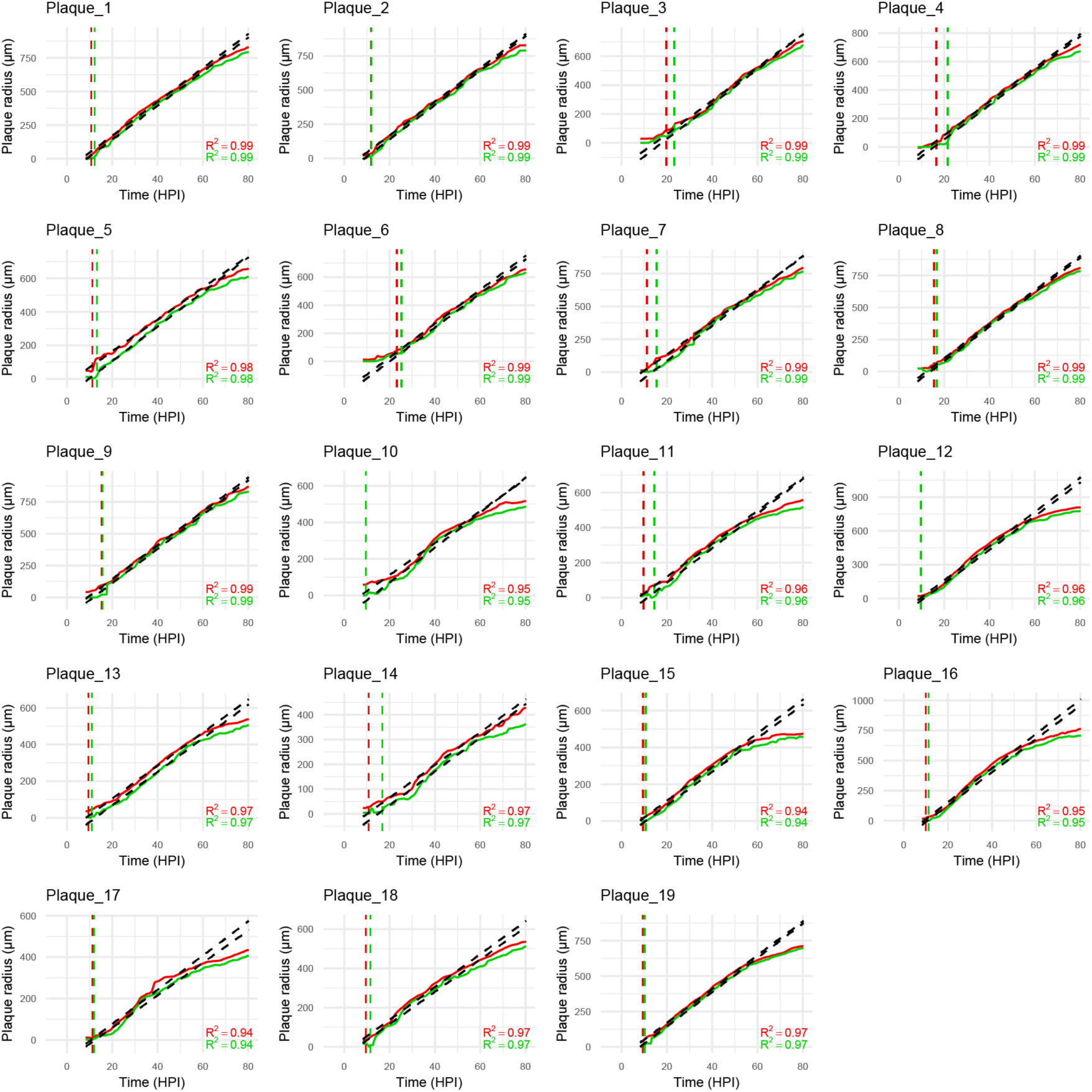
Output of the automated plaque edge detection and analysis pipeline for all plaques analysed. Linear regression of the expansion phase provides the rate of growth, and is indicated by the black dashed lines. The goodness of fit of, and level of variability explained by, the linear model for each plaque and reporter are given by annotated R^2^ value. The transition point from a single to multi-cell infection is identified by the vertical dashed lines. Red indicates the initiation of early gene expression in multiple cells, while green indicates the initiation of late gene expression in multiple cells. These timepoints were termed *t*_0_ and were automatically identified by applying custom filters to the output of a changepoint detection algorithm.

**Supplementary table 1:** Early reporter plaque dynamics for all plaques analysed. Values of *t*_0_ (the initiation of multi-cellular expression) and growth rate for the Early reporter in all plaques analysed.

| Cell line | Plaque number | $t_0$ (HPI) | Growth rate ($\mu\text{m/hr}$) |
| --- | --- | --- | --- |
| **BS-C-1** | 1 | 10.75 | 12.42 |
|  | 2 | 11.75 | 12.18 |
|  | 3 | 19.75 | 11.45 |
|  | 4 | 16.50 | 11.68 |
|  | 5 | 11.25 | 9.29 |
|  | 6 | 23.25 | 11.64 |
|  | 7 | 11.25 | 12.14 |
|  | 8 | 15.50 | 13.06 |
|  | 9 | 15.25 | 13.30 |
|  | **mean** | 15.03 | 12.14 |
| **HaCaT** | 1 | 9.50 | 8.51 |
|  | 2 | 9.75 | 9.04 |
|  | 3 | 9.75 | 14.50 |
|  | 4 | 9.50 | 8.90 |
|  | 5 | 10.75 | 6.45 |
|  | 6 | 9.50 | 8.68 |
|  | 7 | 10.00 | 13.47 |
|  | 8 | 11.25 | 7.77 |
|  | 9 | 9.50 | 8.02 |
|  | 10 | 9.50 | 11.50 |
|  | **mean** | 9.90 | 10.30 |

**Supplementary table 2:** Late reporter plaque dynamics for all plaques analysed. Values of *t*_0_ (the initiation of multi-cellular expression) and growth rate for the Late reporter in all plaques analysed.

| Cell line | Plaque number | $t_0$ (HPI) | Growth rate ($\mu\text{m/hr}$) |
| --- | --- | --- | --- |
| **BS-C-1** | 1 | 12.25 | 12.28 |
|  | 2 | 12.00 | 12.36 |
|  | 3 | 23.25 | 11.99 |
|  | 4 | 21.50 | 11.47 |
|  | 5 | 13.25 | 10.00 |
|  | 6 | 25.25 | 11.76 |
|  | 7 | 15.50 | 12.96 |
|  | 8 | 16.75 | 13.06 |
|  | 9 | 15.75 | 13.38 |
|  | **mean** | 17.28 | 12.59 |
| **HaCaT** | 1 | 9.50 | 9.16 |
|  | 2 | 14.50 | 9.46 |
|  | 3 | 9.75 | 14.26 |
|  | 4 | 11.00 | 9.00 |
|  | 5 | 16.75 | 6.91 |
|  | 6 | 10.75 | 8.59 |
|  | 7 | 11.25 | 13.07 |
|  | 8 | 12.00 | 7.48 |
|  | 9 | 11.50 | 7.74 |
|  | 10 | 10.25 | 11.62 |
|  | mean | 11.73 | 10.44 |

